# Precise Quantification of Translation Inhibition by mRNA Structures that Overlap with the Ribosomal Footprint in N-terminal Coding Sequences

**DOI:** 10.1101/091470

**Authors:** Amin Espah Borujeni, Daniel Cetnar, Iman Farasat, Ashlee Smith, Natasha Lundgren, Howard M Salis

**Affiliations:** Department of Chemical Engineering, The Pennsylvania State University, University Park, PA, 16802; Department of Biological Engineering, The Pennsylvania State University, University Park, PA, 16802

## Abstract

A mRNA’s translation rate is controlled by several sequence determinants, including the presence of RNA structures within the N-terminal regions of its coding sequences. However, the physical rules that govern when such mRNA structures will inhibit translation remain unclear. Here, we introduced systematically designed RNA hairpins into the N-terminal coding region of a reporter protein with steadily increasing distances from the start codon, followed by characterization of their mRNA and expression levels in *E. coli*. We found that the mRNAs’; translation rates were repressed, by up to 1410-fold, when mRNA structures overlapped with the ribosome’s footprint. In contrast, when the mRNA structure was located outside the ribosome’s footprint, translation was repressed by less than 2-fold. By combining our measurements with biophysical modeling, we determined that the ribosomal footprint extends 13 nucleotides into the N-terminal coding region and, when a mRNA structure overlaps or partially overlaps with the ribosomal footprint, the free energy to unfold only the overlapping structure controlled the extent of translation repression. Overall, our results provide precise quantification of the rules governing translation initiation at N-terminal coding regions, improving the predictive design of post-transcriptional regulatory elements that regulate translation rate.

## INTRODUCTION

A mRNA’s translation rate is controlled by its sequence through the collective action of several ribosome-mRNA interactions (1-11), creating a complicated sequence-function relationship. Developing a precise and predictive understanding of these interactions has become essential to controlling protein expression levels for a wide variety of biotechnological applications (12), including biosensor development (13-15), metabolic pathway engineering (16-24), and genetic circuit engineering (25-27). Within this sequence-function relationship, different portions of the mRNA play distinct roles, though most studies have largely focused on how the bacterial ribosome binding site sequence affects the mRNA’s translation rate. Beyond the ribosome binding site, it has been established that a mRNA’s protein coding sequence can greatly affect its translation initiation rate (3,7,28), though the biophysical rules that govern the extent of its control remains unpredictable and poorly quantified. Here, we apply a learn-by-design approach to systematically elucidate the biophysical rules that govern when a protein coding sequence will repress a mRNA’s translation rate, and to precisely quantify the length of the ribosome’s footprint prior to translation initiation.

Translation initiation is a rate-limiting step in gene expression whereby the 30S ribosomal subunit binds to a mRNA’s standby site, hybridizes to its Shine-Dalgarno sequence (SD), and inserts the coding region into its entry channel to form a 30S initiation complex (30SIC), together with tRNA^fMet^ and initiation factors (2,29,30). Afterwards, the 50S ribosomal subunit is recruited to form a 70SIC, GTP is hydrolyzed, and translation elongation begins (31). Notably, the presence of mRNA structures will inhibit 30SIC formation and repress the mRNA’s translation rate, though the magnitude of this effect will depend on the mRNA structures’ locations, thermodynamics, and folding kinetics (8,10,11,32-34). In general, there are three categories of mRNA structures. First, if the mRNA structure is located in the standby site region, upstream of the Shine-Dalgarno sequence, then the ribosome is not required to unfold the structure to initiate translation (10). Instead, a standby site structure will only affect a mRNA’s translation rate by altering the size of the ribosome’s “landing pad”, quantified by the amount of single-stranded RNA available to bind to the ribosome’s platform domain. Second, if a mRNA structure sequesters or overlaps with the Shine-Dalgarno sequence (the 16S ribosomal RNA binding site), then it must be unfolded prior to translation initiation, and therefore the energy needed to unfold those structures will destabilize 30SIC formation (11, 32). Similarly, any mRNA structures that overlap with the spacer or start codon regions also need to be unfolded prior to translation initiation.

However, a third category of mRNA structure exists only within the protein coding region of the mRNA, and it remains unclear when and how such mRNA structures will inhibit translation. The diameter of the ribosome’s Entry channel is about 20 Å wide, and may only accommodate single-stranded RNA, therefore requiring unfolding of the portion of the mRNA that is fed into the channel (29, 35). Further, for the 30S ribosome to bind to the mRNA and form a 30SIC complex, the mRNA that spans its footprint must be entirely single-stranded. Therefore, it is expected that any mRNA structure that overlaps with the 30SIC footprint must be unfolded prior to translation initiation. Several studies have carried out *in vitro* measurements of the ribosome’s footprint, finding that the ribosome extends to between 12 to 19 nucleotides into the mRNA’s N-terminal coding region (36-38). In contrast, the *in vivo* characterization of combinatorial libraries of ribosome binding sites and protein coding sequences has indicated that mRNA structures could inhibit translation when located anywhere from −4 to +37 relative to the start codon (3,7,28). In addition, from an evolutionary conservation perspective, the first 5 to 10 codons (15-30 nucleotides) of a protein coding sequence are biased towards synonymous codons that minimize the formation of mRNA structures (4). More precise *in vivo* measurements of the ribosome’s footprint are needed to accurately predict when an N-terminal mRNA structure will inhibit translation.

In this article, we systematically design a series of synthetic expression cassettes to precisely measure the *in vivo* ribosomal footprint length in *Escherichia coli*. We show that N-terminal mRNA structures will have a dramatic effect on the mRNA’s translation rate, while quantifying the positional and thermodynamic rules that govern their translation repression. By incorporating these improved quantitative rules into our biophysical model of translation initiation, we greatly improve its ability to accurately predict mRNA translation rates across a range of structurally diverse protein coding sequences.

## MATERIAL AND METHODS

### Plasmid Design and Cloning

A series of pFTV1-derived plasmids (ColE1, Cm^R^) (11) were designed and constructed to express a modified mRFP1 fluorescent protein reporter with the objective of introducing rationally designed modifications to the N-terminal region of mRFP1 to introduce specific, desired mRNA structures while preventing the formation of undesired, confounding mRNA structures in non-CDS regions. All plasmid variants utilized a σ^70^ constitutive promoter (BioBrick #J23100) to control *mRFP1* transcription together with a rationally designed ribosome binding site sequence that has the potential to support a high *mRFP1* translation rate in the absence of inhibitory mRNA structures (the no-hairpin control). Specifically, we used the RBS Calculator v2.0 to design a ribosome binding site (RBS) sequence with a 5’ XbaI restriction site, an upstream 6 base pair hairpin, and a 3’ NdeI restriction site with a targeted translation rate of 30000 au on the RBS Calculator’s proportional scale. The resulting 5’ untranslated region (5’-TCTAGAACCCGCCATATACGGCGGGACACACACAAGGAGACCATATG-3’) has an accessible standby site (ΔG_standby_= 0.06 kcal/mol), a high-affinity Shine-Dalgarno sequence (ΔG_SD-antiSD_ = −8.68 kcal/mol), a 3-nucleotide spacer region (ΔG_spacing_ = 1.52 kcal/mol), an AUG start codon (ΔG_start_ = −1.19 kcal/mol), and an overall ribosome binding free energy of ΔG_total_ = −8.3 kcal/mol, yielding a high predicted translation initiation rate of 30400 au. The purpose of the upstream hairpin is to prevent the formation of mRNA structures that sequester the Shine-Dalgarno sequence; the upstream hairpin itself does not overlap with the Shine-Dalgarno sequence and is not predicted to inhibit translation rate. 27 N-terminal mRFP1 variants were designed to incorporate specific, desired mRNA structures, while preventing the formation of alternative mRNA structures. Each variant CDS sequence contains a 5’ NdeI site and a 3’ SacI site.

The starting pFTV plasmid contained a 5’ XbaI upstream of the RBS and a 3’ SacI site inside the N-terminal CDS. To construct the plasmids used in this study, the designed ribosome binding site was first inserted into pFTV1 by annealing two complementary oligonucleotides (Integrated DNA Technologies) with XbaI/SacI overhangs and ligating the insert with digested pFTV1 vector, creating an intermediate plasmid pFTV1a. The CDS variants were then inserted into pFTV1a by constructing DNA fragments with NdeI/SacI overhangs, either by annealing oligonucleotides or by PCR assembly and digestion, followed by ligation with digested pFTV1a. All plasmids were transformed into *Escherichia coli* DH10B cells, followed by sequence verification of isolated clones. All sequences are presented in the **Supplementary Data**.

### Strains, Growth and Characterization

All mRNA level and single-cell fluorescence measurements were performed on plasmid-harboring *Escherichia coli* DH10B cells during long-time cultures, similar to a previous study (10). For each construct, isogenic colonies were used to inoculate overnight cultures in 700 ul LB media supplemented with 50 ug/mL Cm within a 96-well deep-well plate. To begin the characterization, 10 ul culture was diluted into 190 ul of fresh LB/Cm media using a 96-well microtiter plate, and incubated at 37°C with high orbital shaking inside a M1000 spectrophotometer (TECAN). OD_600_ absorbances were recorded every 10 minutes until the OD_600_ reached 0.15, indicating the cells were reaching the mid-exponential phase of growth. At this time, a second 96-well microtiter plate was inoculated by serial dilution using culture from the first plate and fresh LB/Cm media. In the same way, a third serial dilution was conducted using a third 96-well microtiter plate, yielding a total culture time of about 24 hours where cells are continuously maintained in the exponential phase of growth. For each culture, single-cell mRFP1 fluorescence measurements were performed by collecting 10 ul from the end of the second and third dilution, transferring to a microtiter plate with 200 ul PBS solution with 2 mg/ml kanamycin, and utilizing a Fortessa flow cytometer (BD Biosciences) to record 100,000 single-cell fluorescence levels. All single-cell fluorescence distributions were unimodal. The arithmetic mean of distributions is calculated, and the background autofluorescence of *Escherichia coli* DH10B cells is subtracted. All reported fluorescence levels are the average of four measurements from cultures carried out on two separate days, and are listed in **Supplementary Data**.

mRNA level measurements were performed on selected strains by using overnight cultures to inoculate 5 mL Cm-supplemented LB media at 37°C with 300 RPM shaking. Cells were harvested once they reached an OD_600_ absorbance of 1.5 to 2.0, measured using a cuvette-based spectrophotometer (NanoDrop 2000C), and their total RNA extracted using the Total RNA Purification kit (Norgen Biotek), followed by non-specific degradation of contaminant DNA using the Turbo DNAse kit (Ambion). Following extraction, cDNA was prepared using the High Capacity cDNA Reverse Transcription kit (Applied Biosystems). Taqman-based qPCR was performed using an ABI Step One real-time thermocycler (Applied Biosystems), utilizing a Taqman probe targeting a non-modified mRFP1 region (5’-ACCTTCCATACGAACTTT-3’), a forward primer (5’- ACGTTATCAAAGAGTTC-3’), and a reverse primer (5’-CGATTTCGAACTCGTGACCGTTAA-3’). Taqman-based RT-qPCR measurements were also performed on 16S rRNA as an endogenous control, and were used to calculate relative mRNA levels from ΔC_t_ numbers.

### A Biophysical Model of Translation Initiation

We previously developed a statistical thermodynamic model of ribosome-mRNA interactions to predict a mRNA’s translation initiation rate (*r*) from its sequence by calculating the total change in Gibbs free energy (ΔG_total_) when the 30S ribosomal subunit binds to a mRNA at a selected start codon (8,10,11). The ribosome’s binding free energy is related to the mRNA’s translation initiation rate according to Boltzmann’s relationship,

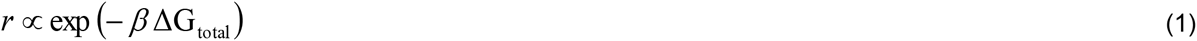

where *β*is the apparent Boltzmann constant, which has been empirically measured to be 0.45±0.05 mol/kcal (8). The total change in Gibbs free energy is calculated using the following multi-term free energy model,

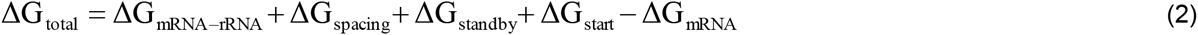

where each term quantifies the strengths of the molecular interactions that control translation initiation in the initial state (free, folded mRNA) and in the after state (a 30S initiation complex) as shown in Figure 1A. ΔG_total_ is the amount of free energy (work) needed to transition the system from the initial to final state. Starting from the initial state, we calculate the amount of free energy needed to fully unfold the mRNA from its minimum-free-energy configuration (ΔG_mRNA_). We then calculate the large amount of free energy that can be released when the 30S ribosomal subunit binds to the mRNA by identifying the mRNA structure and ribosome-mRNA interaction that minimizes the free energy of the final state (ΔG_mRNA-rRNA_ + ΔG_spacing_ + ΔG_standby_ + ΔG_start_), where ΔG_standby_ quantifies the energy needed for the ribosome to bind to the standby site upstream of the Shine-Dalgarno sequence (ΔG_standby_ > 0), ΔG_mRNA-rRNA_ quantifies how much energy is released when the 16S rRNA hybridizes to the mRNA and the mRNA refolds into non-inhibitory structures (ΔG_mRNA-rRNA_ < 0), ΔG_spacing_ quantifies the energetic penalty for stretching or compressing the ribosome, due to non-optimal spacing between the Shine-Dalgarno and start codon (ΔG_spacing_ > 0), and ΔG_start_ quantifies the energy released when the tRNA hybridizes to the start codon. The free energy ΔG_mRNA-rRNA_ can be further broken into the following terms:

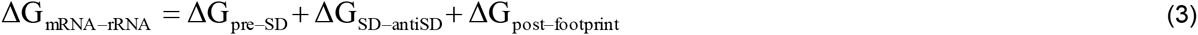

where ΔG_SD-antiSD_ is the free energy released when the last 9 nucleotides of the 16S rRNA hybridize to the mRNA at the Shine-Dalgarno (SD) sequence, ΔG_pre-SD_ is the free energy released when the mRNA upstream of the SD folds into a non-inhibitory structure, and ΔG_post-footprint_ is the free energy released when the CDS portion of the mRNA beyond the ribosome’s footprint region folds into a non-inhibitory structure. We previously developed experimentally validated models for calculating ΔG_standby_ and ΔG_spacing_ (10, 11). All RNA folding energetics and RNA-RNA hybridization free energies are calculated using semi-empirical RNA free energy models (36, 37), provided by the Vienna RNA suite (version 1.8.5) (39). Overall, the biophysical model’s predictions have been experimentally validated by characterizing 495 mRNAs with diverse sequences and structures in several gram-positive and gram-negative bacterial hosts (9-11,15,40), showing that the thermodynamic model can predict 57% of the mRNAs’; translation rates to within 2-fold, and 83% of the mRNAs’; translation rates to within 5-fold, across a 100,000 proportional scale. A key assumption of the thermodynamic model is that there is sufficient time for the mRNA to refold during cycles of translation initiation. In a recent study, we show that RNA folding kinetics can have a significant effect on a mRNA’s translation rate according to a Ribosome Drafting mechanism (8). Here, we use Kinfold to carry out kinetic RNA folding simulations (41) as part of the rational design process to ensure that all studied mRNA structures will rapidly fold to their minimum-free-energy configurations, which will minimize the Ribosome Drafting effect. The biophysical model calculations, Kinfold-calculated RNA folding times, and translation rate predictions for all studied mRNAs are included in **Supplementary Data**.

**Figure 1.**
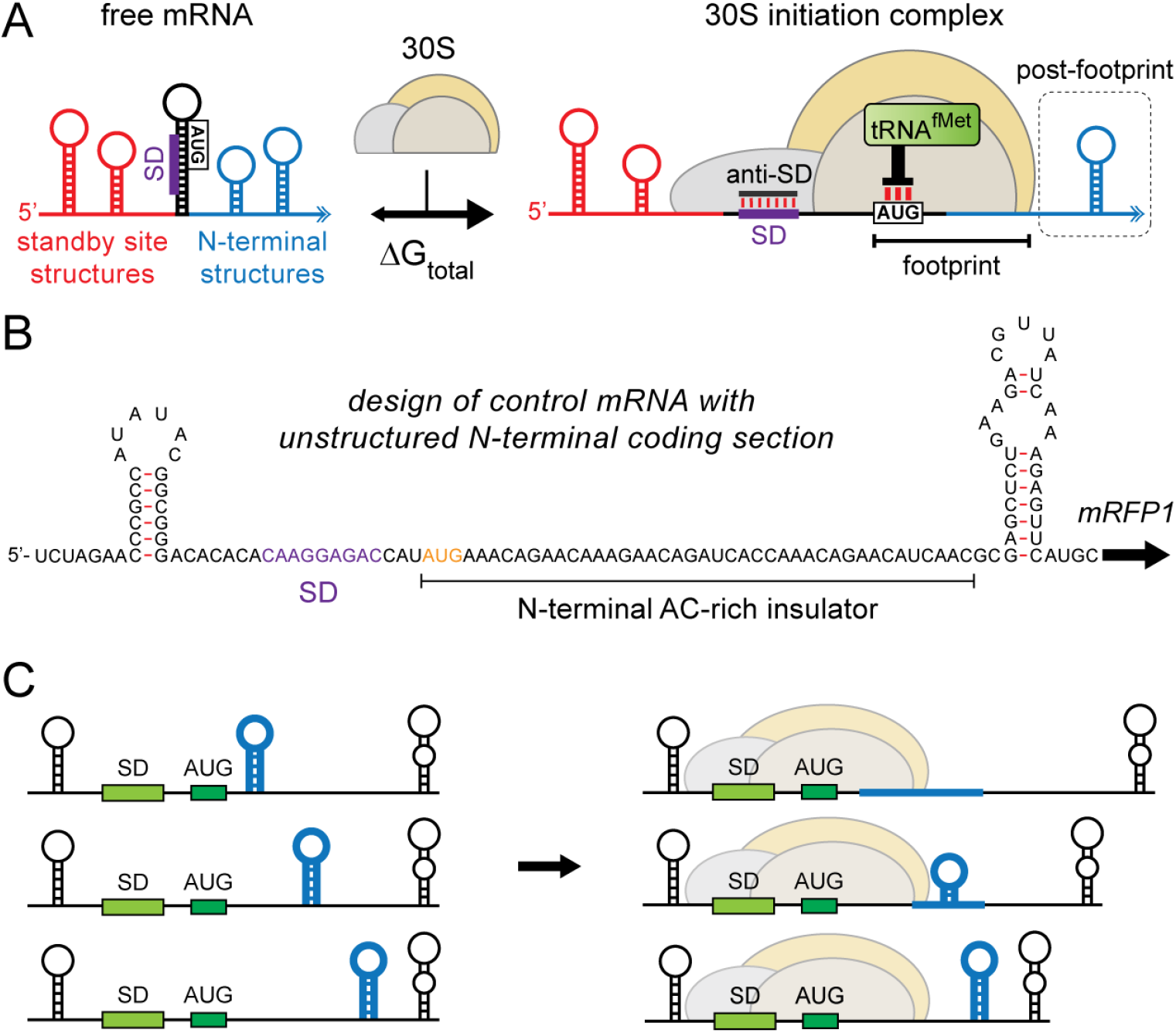
Translation initiation rate is controlled by the overlap between the ribosomal footprint and N-terminal mRNA structures. (A) A biophysical model uses a mRNA’s sequence to calculate the ribosome’s total binding free energy change, which is then related to the mRNA’s translation initiation rate. mRNA structures that overlap with the standby site, with the Shine-Dalgarno sequence, and with the coding sequence have different effects on the ribosome’s binding free energy. (B) A baseline no-hairpin control mRNA contains a highly accessible standby site, an optimized Shine-Dalgarno sequence, and an AC-rich CDS fusion that translates a *mRFP1* reporter at a very high rate, while enabling the insertion of new hairpin-forming CDS fusion sequences without the formation of undesired mRNA structures. (C) Hairpin-forming sequences were inserted into the N-terminal CDS of *mRFP1* at varying distances from the start codon to determine where the mRNA hairpins overlap with the ribosomal footprint.

## RESULTS

### Design of Expression Systems to Measure the Effect of mRNA Hairpins in N-terminal Coding Sequences

We applied a learn-by-design approach to investigate how the location of a stable mRNA hairpin inside the N-terminal of a coding sequence influences a mRNA’s translation initiation rate. Overall, we expected that mRNA hairpins located inside the ribosomal footprint would inhibit the mRNA’s translation rate. More specifically, our sequences were designed to precisely measure the length of the ribosomal footprint in an *in vivo* physiological environment, and to quantify the magnitude of translation repression according to the mRNA hairpins’ locations and folding free energies.

First, we designed, constructed, and characterized a baseline mRNA sequence that enabled us to measure the effect of hairpin-forming sequences within the mRFP1 N-terminal coding sequence, while avoiding the formation of undesired mRNA structures and ensuring that the ribosome’s translation elongation rate does not become a rate-limiting step in the translation process (Figure 1B). The baseline mRNA sequence, called our “no hairpin” control, contains a stable mRNA hairpin upstream of the Shine-Dalgarno sequence, an optimized Shine-Dalgarno sequence that supports a very high translation initiation rate of *mRFP1* (30400 au on the RBS Calculator v2.0 scale), and a 39 nucleotide AC-rich in-frame insertion into the mRFP1 coding sequence. To accelerate translation elongation, the AC-rich CDS fusion utilized codons that are predominantly found in natural, highly translated E. coli coding sequences, called “fast codons”. To minimize any ribosomal pausing, the AC-rich CDS fusion does not contain any Shine-Dalgarno-like sequences. We applied flow cytometry and RT-qPCR to measure the mRFP1 fluorescence and mRNA level of the no-hairpin control in *E. coli* DH10B cells to confirm that it indeed expressed the mRFP1 reporter at a very high level (Figure 2A).

We then designed, constructed, and characterized sets of mRNAs where we inserted hairpin-forming sequences into the mRFP1 N-terminal CDS, systematically varying the location of their 5’ end from +4 to +40 nucleotides after the start codon (Figure 1C). The first set of sequences formed short mRNA hairpins with a 6 base pair duplexed stem. To create in-frame CDS fusions, the hairpins’ loop lengths were varied from 6 to 8 nucleotides. Accordingly, when they are inserted into the N-terminal CDS region at varying positions, their calculated folding free energies ranged from −9.5 to −11 kcal/mol. The second set of sequences formed long mRNA hairpins with a 11 base pair bulged stem and 6 to 8 nt loop lengths. Their calculated folding free energies ranged from −17.2 to −18.6 kcal/mol when inserted at varying positions within the N-terminal *mRFP1* CDS region. When designing these hairpin-forming sequences, we minimized the introduction of confounding variables, such as changes in translation elongation rates and mRNA stability, by ensuring that all hairpin-forming sequences utilized fast codons, and by preventing the appearance of any mRNA duplexes above 8 base pairs to minimize mRNA degradation (42). For each mRNA, we also carried out 1000 kinetic RNA folding simulations using Kinfold (41), and calculated their average folding time, to ensure that all hairpin-forming sequences could rapidly fold to their minimum-free-energy structure. We then characterized the steady-state mRFP1 fluorescence levels and mRNA levels during long-time *E. coli* DH10B cultures maintained in the exponential growth phase to determine how each N-terminal hairpin-forming sequence affected the mRFP1 translation initiation rate and mRNA stability (**Methods**).

**Figure 2.**
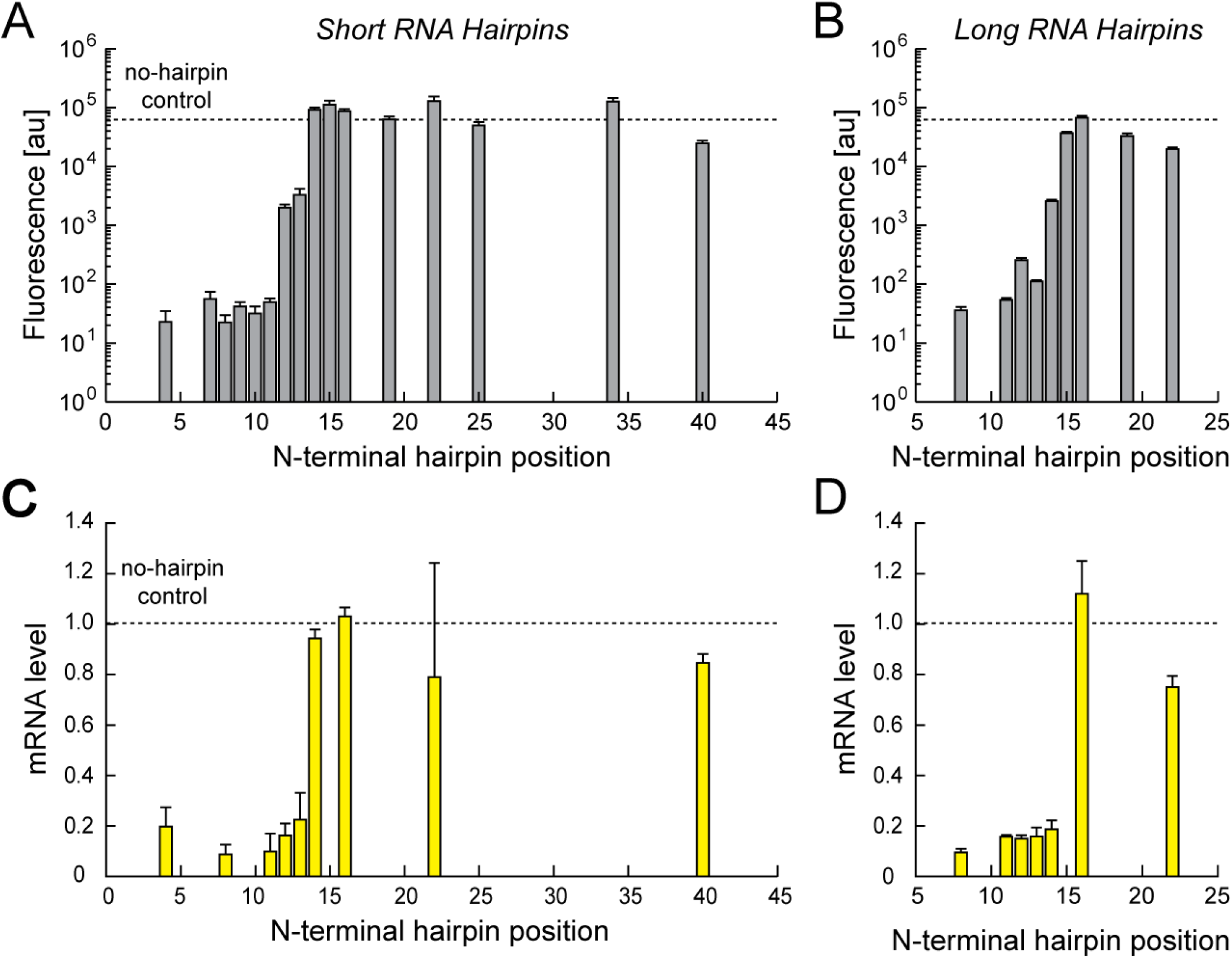
Translation initiation is inhibited based on the position of N-terminal mRNA structures. Measured mRFP1 fluorescence levels when either (A) short mRNA hairpins or (B) long mRNA hairpins were inserted into the mRFP1 coding sequence at designated positions. The mRFP1 fluorescence level of the no-hairpin control is denoted by a dashed line. The measured mRNA levels of selected expression systems with either (C) short hairpins or (D) long hairpins, normalized with respect to the measured mRNA level of the no-hairpin control. Fluorescence data points and error bars are the average and standard deviation of four cytometry measurements on two separate days. mRNA level data points and error bars are the average and standard deviation of 2 to 8 measurements on up to three separate days. All measurements are listed in **Supplementary Data**.

### mRNA Hairpins that Overlap with the Ribosomal Footprint Substantially Inhibit Translation Initiation

Characterization of the designed mRNAs revealed an exceedingly clear and quantitative relationship between the locations of the mRNA hairpins inside the mRFP N-terminus and the mRFP1 expression levels (Figure 2A). In our designed no-hairpin control mRNA that lacks any inhibitory mRNA structures, the mRFP1 fluorescence level was very high (62027.2 ± 6971.4 au) (Figure 2AC, dashed lines). However, when a short mRNA hairpin was introduced downstream of the start codon (position +4), the mRFP1 fluorescence level dropped by 2720-fold to 22.8 ± 12 au (Figure 2A, bars) coincident with a 5-fold drop in mRNA level (Figure 2C, bars), equivalent to a 544-fold decrease in translation rate. Similar magnitudes of translation repression were observed when short mRNA hairpins were positioned at +7, +8, +9, and +10 nucleotides after the start codon. This pattern of translation repression suggests that the ribosome (the 30SIC) was required to unfold the entirety of these mRNA hairpins in order to insert the mRNA into its entry channel, and that the ribosomal footprint extends at least 10 nucleotides into the N-terminal coding sequence.

Next, as the short mRNA hairpin’s position was moved further downstream from the start codon, the magnitude of translation repression smoothly diminished; at positions +11, +12, and +13, the mRFP1 translation rates were repressed by only 122-fold, 5.0-fold, and 4.1-fold, respectively, compared to the no-hairpin control. Because the mRNA’s translation rate underwent a smooth transition from full to partial repression, these measurements suggest that, when a mRNA hairpin only partially overlaps with the ribosomal footprint, the ribosome need only partially unfold the mRNA hairpin. Below, we will detail calculations that explain how partial unfolding of mRNA hairpins requires less free energy, for example, when a portion of the mRNA hairpin can refold into a less energetic structure. Finally, when short mRNA hairpins were positioned from +14 to +40, translation repression was not observed. These measurements show that when a mRNA hairpin does not overlap with the ribosomal footprint, the 30SIC does not need to unfold the hairpin to initiate translation. Instead, the mRNAs’; translation rates were very similar to the no-hairpin control’s translation rate with only minor changes, for example, a 2-fold decrease at position +40 and a 2-fold increase at position +22, that could arise from other interactions. Overall, we observed a 1409-fold change in translation rate as sixteen short mRNA hairpins were inserted from positions +4 to +40 in the N-terminal mRFP1 coding sequence, illustrating the substantial impact of N-terminally located mRNA structures on the ribosome’s ability to initiate translation. Importantly, we did not observe any significant repression of either translation initiation or translation elongation when short mRNA hairpins were inserted at and beyond position +14.

We next investigated whether the insertion of longer, more stable mRNA hairpins into the N-terminal CDS region would similarly influence the mRNA’s translation initiation rate. The measured mRFP1 fluorescences and mRNA levels from nine mRNAs with longer mRNA hairpins, systematically inserted at positions +8 to +22 downstream of the start codon, revealed a similar pattern of translation repression and mRNA level changes (Figure 2BD). A long mRNA hairpin inserted at position +8 dropped the mRFP1 fluorescence level by 1720-fold with a coincident 10.6-fold drop in mRNA level, compared to the no-hairpin control mRNA, yielding a translation repression of 161-fold. When the long mRNA hairpins were positioned further downstream, the amount of translation repression decreased relatively smoothly until, at position +16, the translation rate was no longer repressed, and was similar to the translation rate of the no hairpin control. Overall, changing the positions of these long N-terminally located hairpins altered the mRNA’s protein expression levels by 1861-fold. Similar to the previous set of short mRNA hairpins, when long mRNA hairpins were inserted at +16 and beyond, they no longer caused a significant amount of translation repression.

### Incorporating mRNA Level Measurements to Determine Changes in Translation Rate

Based on our mRFP1 fluorescence and mRNA level measurements, there was a high degree of coupling between a mRNA’s translation rate and its stability (Figure 2). Whenever the mRNA’s translation is greatly repressed by an N-terminally located hairpin, the mRNA’s level was substantially reduced. In contrast, when either short or long mRNA hairpins were inserted +14 or +16 nucleotides downstream of the start codon, the mRNA’s level remained relatively unchanged. These two observations provide support for a previously proposed mechanism whereby actively translated mRNAs, bound by a sufficiently high density of ribosomes, become protected from RNAse activity (43). Quantitative characterization of this relationship remains a topic for a future study. Here, we focused on identifying how the mRNAs’; translation initiation rates were precisely controlled by the positions of the mRNA hairpins and their folding free energies. We calculated the mRNAs’; actual translation rates by dividing the mRFP1 fluorescence measurements by their corresponding measured mRNA levels (Figure 3AB, red circles and lines), providing a clear relationship between the mRNA hairpins’ positions and translation rate changes. By comparison to the no-hairpin control, we also precisely quantified the apparent amount of work (the free energy change) that the ribosome exerted to unfold the inserted mRNA hairpins (Figure 4C).

**Figure 3.**
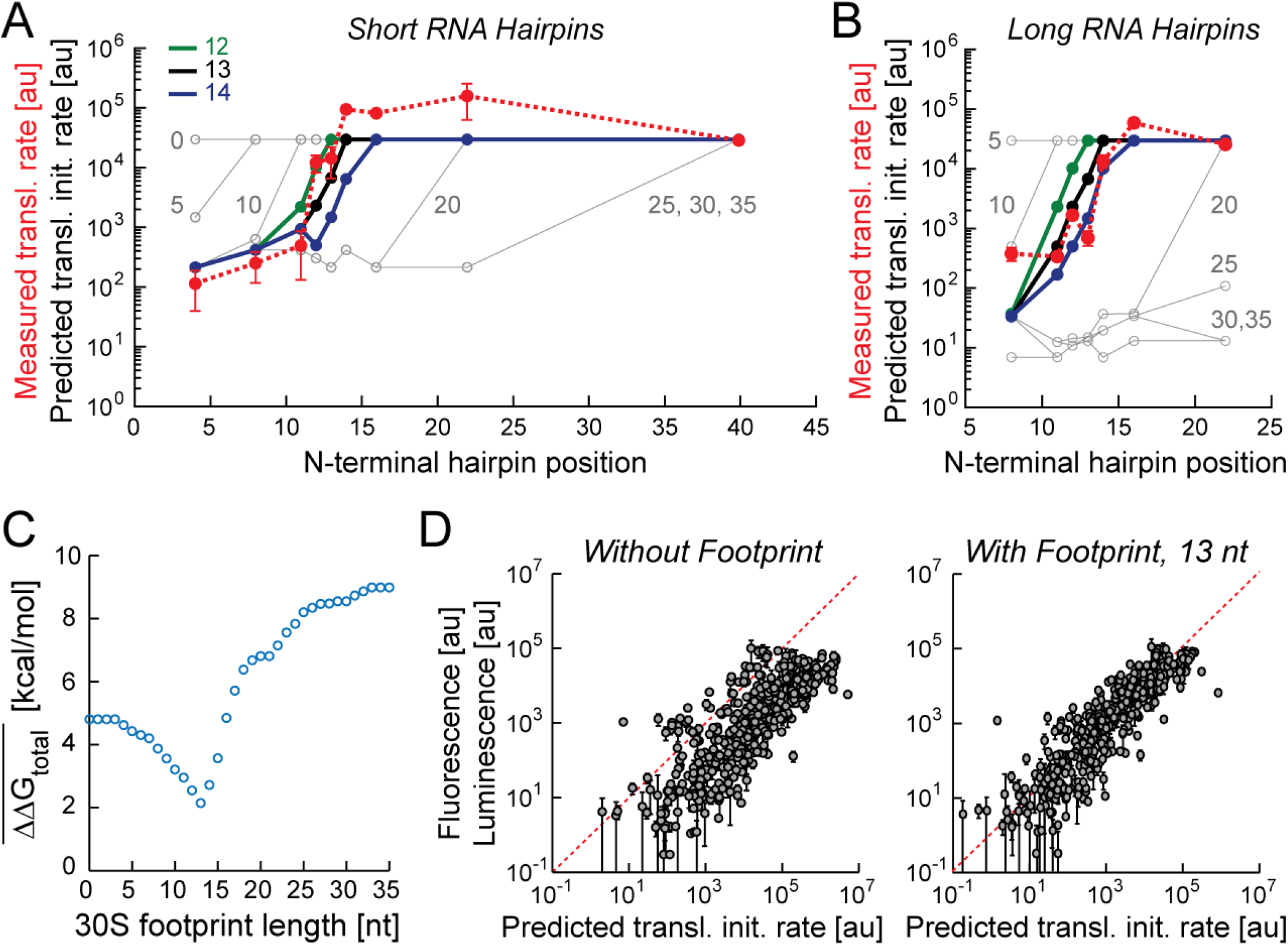
Determination of the Ribosome Footprint Length. Biophysical model predictions using wide range of footprint lengths (0 to 35 nt) are compared to the measured translation rates (red lines, dots) when either (A) a short 6 bp hairpin or (B) a long 11 bp hairpin are located at designated positions in mRFP1’s N-terminal coding section. Model predictions using ribosome footprint lengths of 12, 13, and 14 nt are highlighted (green, black, and blue lines/dots, respectively). (C) A footprint length of 13 nt minimizes the average error in predicted ΔΔG_total_ across 17 characterized mRNAs shown in parts A and B. (D) Incorporation of the ribosome footprint into biophysical model predictions increases its accuracy for 495 previously characterized mRNAs (R^2^ = 0.66, p = 2.5x10^−116^ and R^2^ = 0.78, p = 6.8x10^−164^ for footprint length of 0 and 13 nt, respectively).

### Precise Quantification of the *in vivo* Ribosomal Footprint Length using Biophysical Modeling

Next, by comparing these apparent measurements to our biophysical model’s predictions, we may determine the length of the ribosomal footprint and the characteristics of the mRNA hairpins’ that control the mRNAs’; translation rate. According to our previously developed biophysical model of translation initiation (**Methods**), the length of the ribosomal footprint will have a direct effect on a mRNA’s translation initiation according to a single thermodynamic principle: when the ribosome binds to the mRNA and occupies its lowest free energy state, the portion of the mRNA covered by the ribosome’s footprint must remain unstructured. This principle is implemented by incorporating a single structural constraint for all model predictions. For example, when a mRNA hairpin fully overlaps with the ribosomal footprint, the model determines that the mRNA hairpin must be fully unfolded by the ribosome, resulting in a reduction in the predicted translation initiation rate. In contrast, when a mRNA hairpin does not overlap with the ribosomal footprint, the model determines that the hairpin remains folded in the ribosome-mRNA’s final state, resulting in no change in the predicted translation initiation rate. In between these two cases, when a portion of the mRNA hairpin overlaps with the ribosomal footprint, its effect on the mRNA’s translation initiation rate will depend on the amount of overlap and the possibility of another, less stable mRNA structure forming within the downstream region that is not covered by the ribosome’s footprint.

Specifically, the model calculates the minimum-free-energy of a mRNA region downstream of the start codon, subject to a structural constraint that prevents the formation of mRNA secondary structures within the length of the ribosomal footprint, but allows the refolding of structures outside the ribosomal footprint region. The free energy calculation is designated ΔG_post-footprint_ and is controlled by a single parameter, the ribosomal footprint length. The value of ΔG_post-footprint_ is used to determine ΔG_mRNA-rRNA_, ΔG_total_, and the mRNA’s predicted translation initiation rate according to Equations 1-3. In all model calculations, we do not input the entire mRNA sequence, but instead input the 5’ UTR and a sufficiently large portion of the CDS sequence (at least 75 nucleotides).

We then utilized the developed biophysical model to predict the mRNAs’; translation initiation rates at a range of possible ribosomal footprint lengths to determine the *in vivo* apparent ribosomal footprint length that best reflects our translation rate measurements (Figure 3). We show the mRNAs’; predicted translation initiation rates at a range of putative ribosomal footprint lengths from 0 to 35 nucleotides long and compare these predictions to the mRNAs’; measured translation rates (Figure 3AB, red circles). We carried out this comparison for (Figure 3A) 9 mRNAs with short hairpins and (Figure 3B) 7 mRNAs with long hairpins, where in both cases, we had obtained fluorescence and mRNA level measurements to precisely measure their translation rates. These comparisons sharply identify the *in vivo* ribosomal footprint length to be 12 or 13 nucleotides long for the short hairpin mRNAs and 13 or 14 nucleotides long for the long hairpin mRNAs. Importantly, in both cases, the model correctly predicts the mRNA’s translation rates as the short or long mRNA hairpins’ locations are incrementally shifted. Starting with a hairpin fully overlapping with the ribosomal footprint, and repressing translation, the model is able to predict the sigmoidal increase in translation rate as the mRNA hairpins are shifted into a position that only partially overlaps with the ribosomal footprint. Further shifting of the mRNA hairpin relieves all repression of translation, which is well-predicted by the model.

To further determine the *in vivo* ribosomal footprint length, we combined the 16 characterized mRNAs with long and short hairpins into a single data-set and compared their measured translation rates to the model’s predictions at a range of ribosomal footprint lengths. For each mRNA and ribosomal footprint length, we determined the error in the model’s prediction (ΔΔG_total_) by first using the measured translation rates to determine the apparent change in binding free energy when the ribosome bound to the mRNA (ΔG_total,apparent_), and subtracting the model-calculated ΔG_total,predicted_. We then averaged the absolute value of ΔΔG_total_ across the 16 characterized mRNAs to show the relationship between this average model error versus ribosome footprint length (Figure 3C). Coincident with our visual qualitative analysis in Figure 3AB, we found that the model error reached a global minimum of about 2 kcal/mol when the ribosomal footprint was 13 nucleotides long.

Next, we further examined the accuracy of our identified ribosome footprint length on a much larger data-set of 495 previously characterized mRNA sequences expressing different protein reporters (9-11,15,40). In contrast to the carefully designed mRNA sequences in this study, the large data-set of mRNAs have dissimilar 5’ UTR and protein coding sequences, and their N-terminal coding regions contained mRNA structures with diverse shapes, energies, and positions, providing a stringent test case for the identified ribosome footprint length. We applied the model to predict the translation initiation rates of the 495 mRNAs, using ribosomal footprint lengths of either 0 or 13 nucleotides (**Supplementary Data**), and compared model predictions with their measured expression levels. Here, in the absence of mRNA level measurements, we assumed that the 495 mRNAs’; translation rates were proportional to their measured reporter expression levels. We found that utilizing the correct ribosome footprint length of 13 nucleotides greatly increased the accuracy of biophysical model; the average error in the ΔG_total_ calculation dropped from 6.46 to 2.06 kcal/mol, and consequently increased the accuracy of predicted translation initiation rates (R^2^ = 0.66, p = 2.5x10^−116^ using a footprint length of 0 nt, compared to R^2^ = 0.78, p = 6.8x10^−164^ using a footprint length of 13 nt) (Figure 3D). This analysis shows that a correct measurement of the ribosome’s footprint length is important to accurately predicting the translation initiation rates of a collection of mRNAs with diverse sequences and structures.

**Figure 4.**
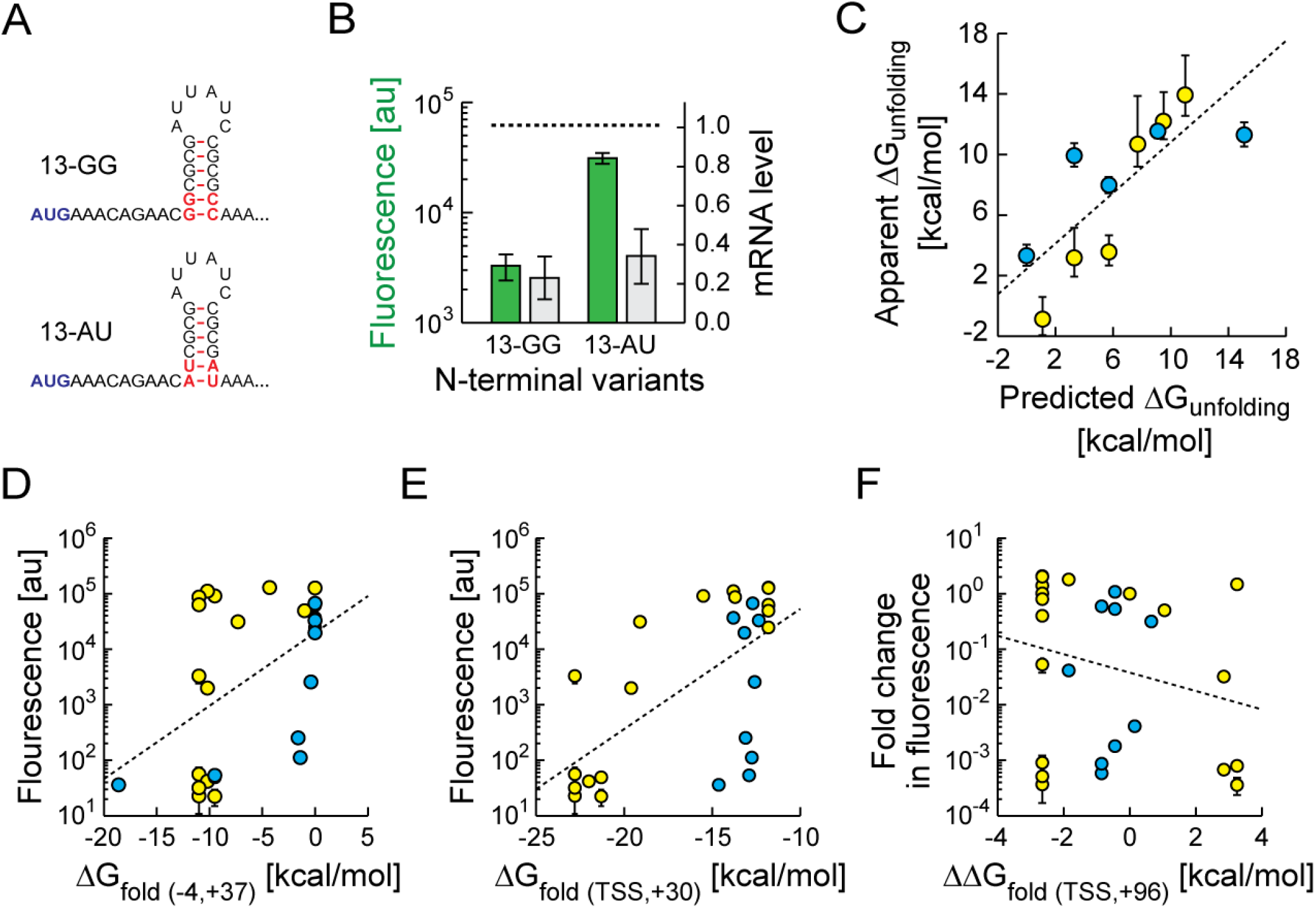
RNA Hairpin Folding Free Energy Controls Translation Inhibition. (A) Two mRNA hairpins were inserted at position +13 with different duplex sequences and folding free energies, but similar hairpin sizes. (B) Fluorescence and mRNA level characterization revealed that the mRNA with the more stable hairpin had a lower translation rate, compared to the mRNA with the less stable hairpin. The horizontal dashed line shows the fluorescence and mRNA level for the no-hairpin control mRNA. (C) The apparent amount of free energy needed to unfold N-terminal mRNA hairpins is quantitatively similar to the model-calculated free energy of hairpin unfolding (R^2^ = 0.62, p = 0.004). Yellow and blue circles are the short and long RNA hairpins, respectively. (D and E) The folding free energies of the mRNA regions spanning either (D) from −4 to +37 or (E) from the transcriptional start site (TSS) to +30 correlate poorly with their measured fluorescence levels (R^2^ = 0.24, p = 0.01; and R^2^ = 0.41, p = 3.1x10^−4^, respectively). +1 is the start codon. (F) The relative changes in the mRNA’s expression level (fluorescence level) did not correlate to the differences in folding free energy for the mRNA region spanning from TSS to +96, compared to the no-hairpin control mRNA (R^2^ = 0.06, p = 0.23). Parts D, E, and F show the measured fluorescence levels for all 27 characterized mRNAs in this study.

### N-terminal RNA Hairpins Inhibit Translation Initiation According to their Folding Free Energy

We next investigated the relationship between the RNA structures’ folding free energies and the mRNAs’; translation initiation rates, and how the specific details of the RNA folding calculation will influence this relationship. Several previous studies have observed that reducing the thermodynamic stability of mRNA structures near the beginning of a coding sequence will have the overall qualitative effect of increasing a mRNA’s translation rate (3,7,28), though in each study, the thermodynamic calculations were performed on different mRNA regions arriving at different types of correlations. For example, in Kudla et. al., the highest correlation between the RNA folding free energy and measured fluorescence levels was observed when the folding calculation was performed specifically on the mRNA region spanning −4 to +37 (+1 is the start codon beginning). Similar analyses were performed in Kosuri et. al. and Goodman et. al., where they found the folding free energies of the mRNA regions TSS to +30 or TSS to +96, respectively, had the highest correlation to the mRNAs’; translation rates (TSS is the transcriptional start site and +1 is the start codon beginning). Importantly, the characterized mRNAs in Kudla et. al. and Goodman et. al. had variable N-terminal coding sequences and constant 5’ untranslated regions, which are similar to the designed mRNAs characterized in this study. However, based on our measurements here, translation initiation is only inhibited by the presence of an N-terminal mRNA structure when it overlaps with the ribosomal footprint, which is located from +1 to +13. Performing a folding calculation on a much longer mRNA region, which includes both inhibitory and non-inhibitory mRNA structures, may not yield a reproducible correlative relationship on different sets of mRNAs. Therefore, we revisited how these various approaches to the folding calculation affected the observed relationship between RNA folding free energy and measured translation rate.

We first tested the sensitivity of the relationship between RNA unfolding free energy and mRNA translation rate, considering only inhibitory RNA hairpins. To do this, we designed and characterized a new mRNA variant that contains a slightly weakened RNA hairpin positioned exactly at +13 (Figure 4A). We swapped the terminal dinucleotide duplex of the hairpin from GG:CC to UA:AU, changing the hairpin’s unfolding free energy from 11 to 7.3 kcal/mol, and thereby making it easier for the ribosome to unfold the dinucleotide overlapping with the ribosomal footprint at position +13. We then measured the fluorescence and mRNA level of the modified mRFP1-expressing mRNA, and found that the mRFP1 reporter expression level increased by 9.5-fold, while the mRNA level marginally increased by 1.5-fold, resulting in a 6.2-fold increase in measured translation rate (Figure 4B). Consistent with this measurement, the biophysical model predicts that weakening the RNA hairpin should have increased the mRNA’s translation rate by 5.3-fold. This data provides an additional demonstrative example showing that, when a mRNA structure overlaps with the ribosomal footprint, its unfolding free energy has a large effect on a mRNA’s translation rate.

Next, we tested the quantitative relationship between the apparent and predicted folding free energies of inhibitory mRNA structures. We utilized our collection of characterized mRNAs that contain long or short hairpins that fully overlap with the ribosomal footprint and therefore must be fully unfolded by the ribosome to initiate translation. From the translation rate measurements, we calculated the apparent amount of free energy needed to unfold the inhibitory hairpins, using the following equation:

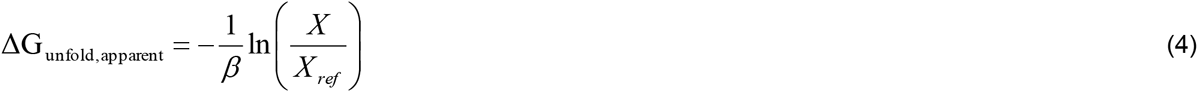

where *X* is the measured translation rate of a mRNA with an N-terminal hairpin that overlaps with the ribosomal footprint, and *X_ref_* is the measured translation rate of the no-hairpin control mRNA that does not have any N-terminal inhibitory hairpins. The constant *β* has been empirically determined to be 0.45 mol/kcal. In Figure 4C, we compared the hairpins’ apparent unfolding free energies to the model’s predictions, and found quantitative agreement (R^2^ = 0.62, p = 0.004), demonstrating that the inhibitory hairpins’ unfolding free energies controlled the mRNAs’; translation rates.

In contrast, when the RNA folding calculation includes both inhibitory and non-inhibitory mRNA structures, we do not observe a quantitative relationship between the mRNA’s folding free energy and the mRNA’s translation rate. The previously proposed RNA folding calculations, spanning −4 to +37, TSS to +30, or TSS to +96, did not result in clear quantitative relationships (R^2^ = 0.24 for [-4, +37], R^2^ = 0.41 for [TSS, +30], and R^2^ = 0.06 for [TSS, +96]) (Figure 4DEF). The RNA folding calculations from −4 to +37 or from TSS to +30 will often exclude N-terminal mRNA structures that partially overlap with the ribosomal footprint, while the calculation from TSS to +96 will include both inhibitory and non-inhibitory mRNA structures. These results show that utilizing any constant cutoff in the mRNA folding calculation may exclude the mRNA structures that inhibit translation, or include mRNA structures that have no effect on translation. Instead, the correct RNA folding calculation should only consider mRNA structures that overlap or partially overlap with the ribosomal footprint.

## DISCUSSION

In this work, we applied a reductive learn-by-design approach to precisely determine the N-terminal mRNA structures that need to be unfolded by the bacterial ribosome during translation initiation. We designed 27 mRNAs with different N-terminal coding sequences, systematically varying the positioning and energetics of their structures, followed by characterization of their translation rates in *E. coli* by combining fluorescent protein and mRNA level measurements (Figure 1). We found that protein expression levels were repressed, by up to 5800-fold, when short or long mRNA structures overlapped with the *in vivo* ribosomal footprint (Figure 2AB). When mRNA level measurements were taken into account, the mRNAs’; apparent translation rates were repressed by up to 1410-fold under the same conditions (Figure 2CD). In contrast, when the mRNA structure was located outside the ribosome’s footprint, protein expression and translation rate were repressed by less than 2-fold. By combining our measurements with biophysical modeling, we precisely determined that the ribosomal footprint extends 13 nucleotides past the start codon (Figure 3ABC). By utilizing this improved quantification of the ribosomal footprint length, we showed that our biophysical model could more accurately predict the translation rates of a collection of 495 characterized mRNAs with diverse sequences and structures (Figure 3D). Finally, we determined that the folding energetics of the N-terminal mRNA structures control the mRNAs’; translation rates, but only when the N-terminal mRNA structure overlaps with the ribosomal footprint (Figure 4ABC). Our maximally informative measurements and biophysical modeling calculations are an improvement over previous “big data” studies where correlations between translation rate and various RNA folding energy calculations were observed, but not tested for mechanistic causality (Figure 4DEF).

Our measured value of the *in vivo* ribosomal footprint length is consistent with previous studies that have utilized a variety of different techniques (36-38). First, Hüttenhofer and Noller applied chemical footprinting on *in vitro* mRNA-30S ribosome complexes to measure the extent of protection from hydrolysis, and found that the ribosomal footprint extended to +19 nucleotides past the start codon when initiator tRNA^fMet^ was added, but was shortened to +5 nucleotides when tRNA^fMet^ was absent (38). Interestingly, inserting a stable mRNA structure from position +10 to +21 resulted in a loss of protection past +5 within the ribosome-mRNA complex. Second, by adding regulatory small RNAs that bind to the N-terminal coding sequence, Bouvier et. al. demonstrated that small RNAs can repress translation initiation if their binding site overlaps with the ribosomal footprint, which was measured to be 14 ± 2 nucleotides, depending on the small RNA, referred more generally as the first five codon rule (37). By conducting *in vitro* toeprinting of ribosome-mRNA complexes with antisense DNA oligos, they also found that 30S ribosome-mRNA complex formation was strongly inhibited when DNA oligos covered the mRNA at or before position +12. Third, by applying optical tweezers to monitor the ribosome-catalyzed unfolding of mRNA structures, Qu et. al. found that the ribosomal footprint was 12 ± 2 nucleotides, and that during translation elongation, mRNA structures were unwound through a combination of biased thermal ratcheting and mechanical opening (36). Interestingly, the ribosome’s ability to mechanically open mRNA structures, utilizing GTP hydrolysis, ensures a minimum rate of translation elongation regardless of the mRNA structure’s stability. However, prior to translation initiation, GTP hydrolysis does not take place and therefore the ribosome relies on biased thermal ratcheting to unwind mRNA structures. This distinction explains why the folding free energies of mRNA structures within the N-terminal coding sequence have a significant effect on a mRNA’s translation initiation rate, but not its translation elongation rate.

Overall, our integrated computational design and experimental approach enabled us to elucidate and quantify the physical rules that govern when the ribosome unfolds N-terminal mRNA structures inside cells and how their unfolding energetics controls the mRNA’s translation initiation rate. The quantification of these rules improved our ability to predict a mRNA’s translation initiation rate according to its sequence, and thereby accelerates the rational design of mRNAs, riboswitches, and other post-transcriptional regulatory elements that manipulate translation initiation rates for useful purposes (15,25,40).

## SUPPLEMENTARY DATA

Supplementary Data are available at NAR online.

## FUNDING

This research was supported by the Air Force Office of Scientific Research (FA9550-14-1-0089), the Office of Naval Research (N00014-13-1-0074), and an NSF Career Award (CBET-1253641) to H.M.S.

## CONTRIBUTIONS

A.E.B. and H.M.S designed the study, analyzed results, and wrote the manuscript. A.E.B. and H.M.S developed the model. A.E.B., D.P.C. I.F., A.S., and N.L. conducted the experiments.

### Conflict of interest statement

None.

